# Adaptive divergence and the evolution of hybrid trait mismatch in threespine stickleback

**DOI:** 10.1101/2020.11.26.400440

**Authors:** Avneet K. Chhina, Ken A. Thompson, Dolph Schluter

## Abstract

Hybrid fitness can be negatively impacted by trait mismatch, whereby hybrids resemble one parent population for some phenotypic traits and the other parent population for other traits. In this study, we used threespine stickleback fish (*Gasterosteus aculeatus* L.) to test whether trait mismatch in hybrids increases with the magnitude of phenotypic divergence between parent populations. We measured morphological traits in parents and hybrids in crosses between a marine population representing the ancestral form and twelve freshwater populations that have diverged from this ancestral state to varying degrees according to their environments. We found that trait mismatch was greater in more divergent crosses for both F_1_ and F_2_ hybrids. In the F_1_, the divergence–mismatch relationship was caused by traits having dominance in different directions whereas it was caused by segregating phenotypic variation in the F_2_. Selection against mismatched traits is an ecological analogue to selection against intrinsic hybrid incompatibilities, and our results imply that extrinsic hybrid incompatibilities accumulate predictably as phenotypic divergence proceeds.

## Introduction

The evolution of reduced hybrid fitness—post-zygotic isolation—is a crucial component of the speciation process (Coyne and Orr, 2004). Post-zygotic isolation is often associated with the buildup of intrinsic genetic incompatibilities that accumulate as populations adapt and diverge (Coughlan and Matute, 2020). Yet, many young species lack strong genetic incompatibilities and hybrid fitness is instead determined by how the phenotype of hybrids facilitates their interactions with complex ecological environments to influence performance (Arnold, 1983). Hybrids can have poor fitness if they have an intermediate phenotype in an environment where there is no intermediate niche (e.g., distinct host plants; Bendall et al. 2017; Matsubayashi et al. 2010; Zhang et al. 2021). Alternatively, if hybrids possess mismatched combinations of traits from both parent populations, this could reduce their fitness even where intermediate environments do exist (Arnegard et al., 2014; Thompson et al., 2021). If the extent of trait mismatch increases over the course of divergence between populations in a manner similar to intrinsic genetic incompatibilities (Coyne and Orr, 1989, 1997; Edmands, 1999), this would suggest that ecologically-mediated natural and/or sexual selection against hybrids could decline predictably as parent populations diverge.

Trait mismatch results from genetic dominance and/or the segregation of divergent alleles in recombinant hybrids. Dominance can cause mismatch in hybrids if some traits are dominant toward one parent and other traits are dominant toward the other, and a recent synthesis study suggests that this is common in F_1_ hybrids (Thompson et al., 2021). Because segregation and recombination in the F_1_ generation weakens trait correlations, individual back-cross or F_2_ hybrids will, by chance, inherit alleles that result in the expression of trait combinations not seen in parents (Schemske and Bradshaw, 2002). Because dominance affects the F_1_ more than the F_2_ (Lynch and Walsh, 1998), mismatch in F_1_s is primarily expected to be a result of dominance whereas mismatch in recombinant hybrids might be due to one or both of dominance and segregation variance.

First principles and theory predict that the magnitude of trait mismatch in hybrids should be positively associated with divergence between parent populations. Mismatch can be defined geometrically for a pair of traits as the average of the distance between individual hybrid phenotypes and a line connecting parental mean phenotypes (Fig. 1A). Dominance alone can cause a divergence-mismatch association (Fig. 1B). Imagine a case where a hybrid expresses one parent population’s trait value for one trait, and the other parent population’s trait value for a second trait. If the parents differ little for these traits, the hybrid will only have a tiny amount of mismatch. If the parents differ substantially, however, the same amount of dominance will generate a greater magnitude of mismatch. The second reason underlying a divergence-mismatch relationship is that the amount of phenotypic variation in the traits of recombinant hybrids—the segregation variance—is expected to increase with the magnitude of phenotypic divergence between parent populations (Barton 2001; Chevin et al. 2014; Slatkin and Lande 1994) (Fig. 1C). Greater segregation variance results in more extreme mismatched trait combinations appearing in hybrids, and therefore this mechanism is expected to generate greater mismatch as the magnitude of divergence between parents increases. Thus, dominance would be expected to underlie a divergence-mismatch relationship in F_1_s, whereas both dominance and segregation variance could cause such a pattern in recombinant hybrids.

**Fig. 1.**
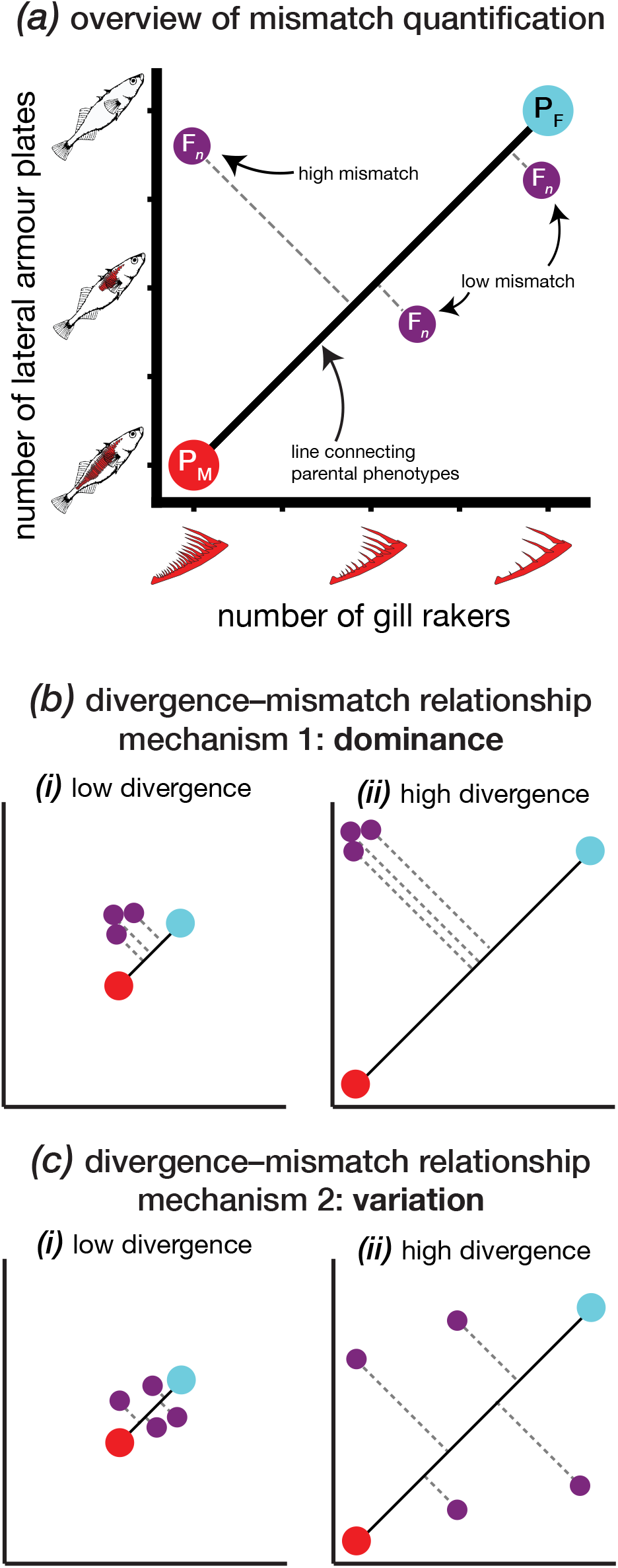
Visual overview of mismatch for two traits. Main text analyses consider multivariate space, but only two traits are shown here for ease of visualization. All panels show parent mean phenotypes as large red (marine parent; P_M_) and blue points (freshwater parent; P_F_) and individual hybrids (F_*n*_) as smaller purple points. Panel (A) shows three individual hybrids—one with high mismatch and two with low mismatch. The length of the dashed lines is the ‘mismatch’ quantity. Panels (B) and (C) show the two mechanisms linking mismatch to phenotypic divergence—dominance and variation. Axes and colours are the same as in panel (A) but labels are omitted for clarity. Both mismatch–divergence panel shows (i) a case of low phenotypic divergence between parents and (ii) a case of high phenotypic divergence between parents.

In this article, we use threespine stickleback fish to test the prediction that trait mismatch in hybrids increases with the magnitude of morphological divergence between parent populations. Freshwater stickleback populations have independently diverged to varying degrees from a common marine ancestor since the last glacial maximum (approx. 10 kya). Contemporary marine populations remain abundant in the sea today and are readily crossed with derived forms. Variation among freshwater populations occurs primarily along a limnetic (i.e., zooplanktivorous) to benthic (i.e., consuming large macro-invertebrates living among the vegetation or lake sediments) axis (Bell and Foster 1994). Although all have adapted to the freshwater habitat, the more limnetic freshwater populations tend to be phenotypically similar to marine populations whereas the more benthic populations are dissimilar. Because more benthic populations have undergone more phenotypic divergence from the marine ancestor, we hypothesized that their hybrids (in crosses with an extant marine population) will exhibit greater mismatch than those produced from crosses with less divergent populations. To test this hypothesis, we measured morphological traits in hybrids from crosses between the ancestral marine form and 12 derived freshwater populations, quantified mismatch, and investigated its causes.

## Methods

### Study system

The threespine stickleback is a teleost fish species distributed throughout the coastal areas of the northern hemisphere (Bell and Foster 1994). Marine stickleback colonized an array of post-glacial lakes and have rapidly adapted to prevailing ecological conditions (Schluter 1996). Stickleback that live in lakes containing predators and other competitor fish species (e.g., prickly sculpin) remain similar to the marine population for many morphological traits (Ingram et al., 2012; Miller et al., 2019). By contrast, populations that have evolved in small lakes with few or no predators and competitors often have more derived phenotypes specialized for foraging on large benthic invertebrates. Three lakes contain ‘species pairs’ with reproductively isolated limnetic and benthic populations (McPhail, 1984, 1992; Schluter and McPhail, 1992)—the limnetics resemble the marine ancestor for many traits, whereas the benthics are among the most derived.

Because adaptive divergence between marine and freshwater populations occurred recently, populations can be readily crossed and typically have few if any ‘intrinsic’ incompatibilities (Hatfield and Schluter 1999; Lackey and Boughman 2017; Rogers et al. 2012). Extant marine populations, within a particular geographic location, are phenotypically similar to the ancestral populations that founded present-day freshwater populations (Morris et al. 2018). We leveraged this continuum of phenotypic divergence using crosses to test the prediction that hybrid mismatch will be greater when more divergent benthos-feeding populations are crossed with a marine ancestral population than when this ancestral population is crossed with less divergent zooplanktivorous populations.

### Fish collection and husbandry

Wild fish were collected in British Columbia, Canada, in April–June of 2017 and 2018. We sampled twelve freshwater populations from nine lakes (Fig. 2A; three lakes [Paxton, Priest, and Little Quarry] contain reproductively isolated benthic-limnetic ‘species pairs’ [McPhail 1992] and thus contributed two populations each). The marine population was collected from the Little Campbell River (Fig. 2A). Fish from the Little Campbell River are anadromous—living in the sea and breeding in freshwater, though we refer to them as ‘marine’ here for consistency with previous studies (e.g., Schluter et al. 2021). Wild fish were caught using minnow traps or dip nets. We crossed six gravid marine females with six males from each freshwater population to generate six unique F_1_ hybrid families per population, and also generated four to six non-hybrid (i.e., ‘pure’) families for each freshwater parental population and the marine ancestor. All offspring were raised in the lab under common conditions (see **Supplementary Methods**). Crosses were conducted in only one direction (marine as dam) to standardize cytoplasm among hybrid crosses and also because obtaining a sufficient number of wild gravid females for some populations was prohibitively difficult. When lab-raised fish reached reproductive maturity, F_1_ hybrids from unrelated families were crossed to make three F_2_ families within each cross population (with the exception of Paxton Lake benthics which, due to aquarium space constraints in 2018, had only two F_2_ families from the same two F_1_ parent families).

**Fig. 2.**
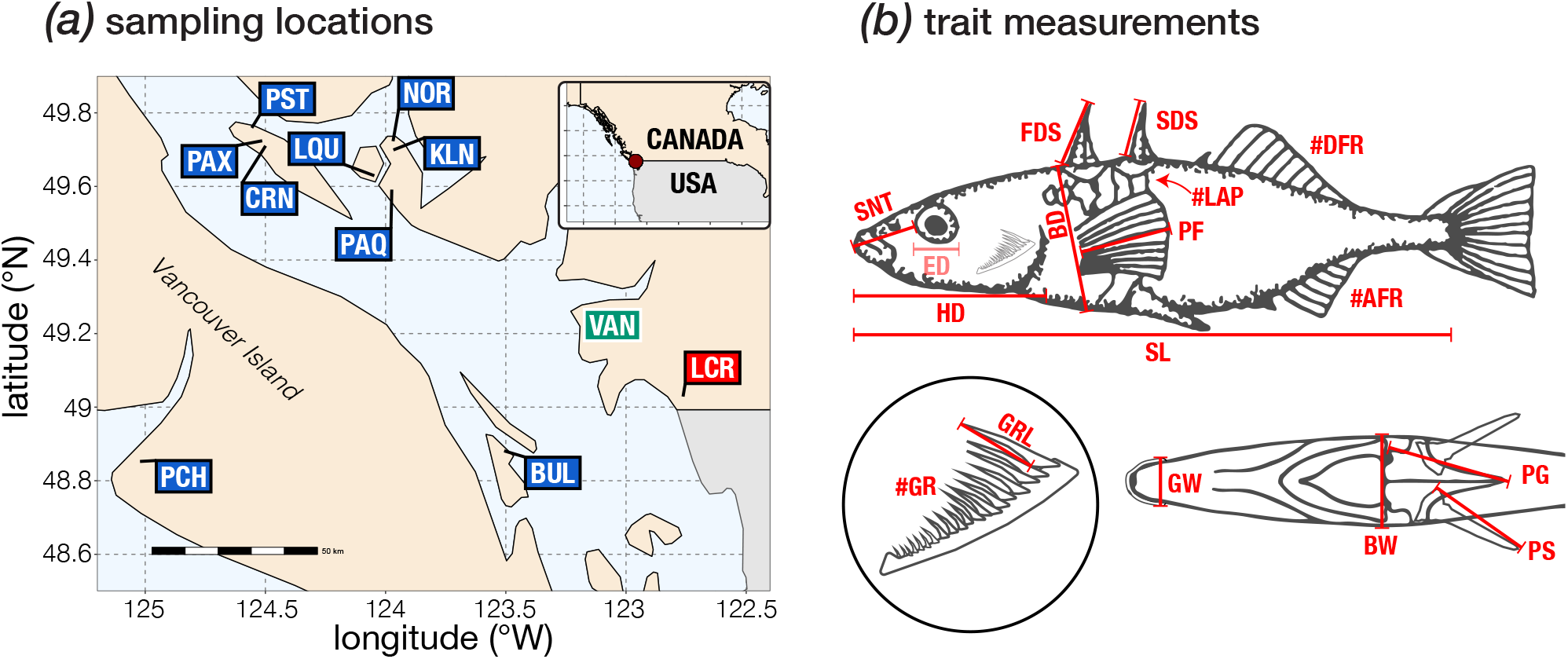
Overview of sampling locations and trait measurements. Panel (a) shows locations where the source populations were collected in British Columbia, Canada. Boxes show collection locations of the marine population (red box; LCR—Little Campbell River) and freshwater populations (blue boxes; left to right: PCH—Pachena Lake; PAX—Paxton Lake; CRN—Cranby Lake; PST—Priest Lake; LQU—Little Quarry Lake; PAQ—Paq (Lily) Lake; NOR—North Lake; KLN—Klein Lake; BUL—Bullock Lake). The green label indicates the location of Vancouver (VAN). Panel (b) shows the measurements of all 16 traits in the dataset and standard length. The upper section of the panel shows the lateral view (traits left to right: SNT—snout length; ED—eye diameter [the transparent shade of red indicates this trait was measured but not analyzed—see ‘Repeatability’ section of methods]; HD—head length; FDS—length of first dorsal spine; BD—body depth; SL—standard length; SDS—length of second dorsal spine; PF— pectoral fin length; #LAP—number of lateral armour plates; #DFR—number of dorsal fin rays; #AFR—number of anal fin rays). The bottom left section of the panel shows a zoomed in drawing of the upper arm of the outer gill raker arch (#GR—number of gill rakers; GRL—length of longest gill raker). The lower right section shows an anteroventral view of the body (GW—gape width; BW—body width; PG—length of pelvic girdle; PS—length of pelvic spine). The upper drawing was originally published by Bell and Foster (1994) and is re-used with permission from M. Bell.

Fish were lethally sampled when individuals in the tank reached a mean standard length of approximately 40 mm. For F_1_s, tanks were sub-sampled and remaining individuals were raised to produce F_2_s. For F_2_s, entire tanks were lethally sampled. Fish had not reached reproductive maturity at the time of sampling, and we therefore could not determine their sex. Fish were preserved in formalin, stained with alizarin red, and then stored permanently in 40% isopropyl alcohol.

### Phenotype measurements

We measured 16 traits and standard length on stained fish (Fig. 2B). For all traits, we measured at least 100 pure marine parents, and 30 pure freshwater parents, 30 F_1_ hybrids, and 60 F_2_ hybrids from each population and marine-freshwater cross (all lab-raised; see summary dataset [**to be archived on Dryad**] for trait means, standard deviations (SDs), and sample sizes for all populations). We used a dissecting microscope to count the number of dorsal fin rays, anal fin rays, lateral armour plates, and gill rakers. We also measured the length of the longest gill raker using an ocular micrometer. We photographed the left and ventral sides of each fish with a Nikon D300 camera and used ImageJ (Abramoff et al. 2004) to make linear measurements of body dimensions and bones. Pectoral fins were dissected, immersed in a more concentrated alizarin red stain for at least 24 hr, then photographed. We measured the length of the longest fin ray as pectoral fin length. All measurements with the exception of eye diameter were highly repeatable (*r* ≥ 0.9; see Fig S1), and as a result all traits except eye diameter were used for subsequent analysis. A small number (*n* = 9) of fish had missing second dorsal spines, which caused them to be extreme outliers. We excluded these fish from the analysis. One additional fish that failed to inflate its swim bladder was also excluded.

We size-corrected all linear measurements by replacing raw measurements with the residuals from simple log-log (ln-transformation) linear regressions with standard length conducted across the entire dataset. We controlled for as much of the variation as possible among populations by sampling them at a relatively consistent mean size. Log-transformation of linear measurements renders trait variances comparable across fish of different sizes (Hatfield, 1997). Some measurements are affected if fish are fixed with an open gape, so we further corrected for fixation position by assigning all fish a number (0, 1, or 2) depending on the extent to which the mouth was open and then performing a further correction as above using residuals for gape width, snout length, and head length. Trait measurements for missing spines (first dorsal spine or pelvic spine) or pelvic girdle were given a raw value of 0.1 mm before log-transformation (the log of 0 is undefined). Unlike the second dorsal spine, variation in the presence of these traits is common and does not result in extreme outliers. Following size-correction, traits were standardized across the entire dataset to a mean of 0 and a standard deviation of 1.

### Data analysis

We first investigated whether trait mismatch was associated with the magnitude of phenotypic divergence between parent populations. Following this, we quantify the role of dominance and trait variation in driving this relationship.

#### Software

All data processing and model-fitting was done using R (R Core Team 2019) using the `tidyverse` (Wickham 2017). Mixed models were fit using `lme4` (Bates et al. 2014) and analysed using `lmerTest` (Kuznetsova et al. 2014 with the Kenward-Roger approximation for the denominator degrees of freedom (Kenward and Roger 1997). The ‘map’ function in `purrr` (Henry and Wickham 2019), and associated functions in `broom` (Robinson et al. 2020), were used to streamline code for iterating models over grouping variables. Partial residuals were plotted using `visreg` (Breheny and Burchett 2017). `for loop` code was streamlined with the functions in `magicfor` Makiyama 2016). We used the `emmeans` package (Lenth et al. 2020) and the ‘cld’ function in `multcomp` (Hothorn et al. 2008) to assist with post-hoc comparisons. The functions in the ‘correlation’ package (Makowski et al. 2019) produced correlation matrices.

#### Quantifying phenotypic divergence

We quantified the magnitude of phenotypic divergence between pure marine and freshwater populations as our main predictor of mismatch. To do this, we calculated the Euclidean distance between each freshwater population’s mean phenotype for all standardized traits and the marine mean phenotype for all traits. For all estimates of population mean phenotypes we use the unweighted mean of family means (we note that our conclusions are unchanged if we average across individuals rather than families).

#### Trait mismatch

Trait mismatch is a quantitative metric capturing the extent to which individual hybrids deviate from the line connecting parental mean phenotypes (Fig. 1; Thompson et al. 2021). Mismatch is a response variable in our analyses and is measured for individual hybrids. We consider all traits together in multivariate space. Correlations between pairs of traits in F_2_ hybrids were low (median |*r*_Pearson_| = 0.2), and most (87 %) were not statistically significant at *P* = 0.05, and for this reason we retain original traits and do not use dimensionality-reduction techniques such as principle components analysis. (Conclusions from a separate set of analyses considering pairs of traits at a time [i.e., ‘pairwise’ mismatch *sensu* Thompson et al. 2021] were similar to those from the multivariate analysis and for simplicity are shown only in the Supplementary Results.)

Mismatch is the shortest (i.e., perpendicular) Euclidean distance between a hybrid’s phenotype and the line that connects the two parental mean phenotypes (Fig. 1). Mismatch (*d*_mm_) was calculated as:

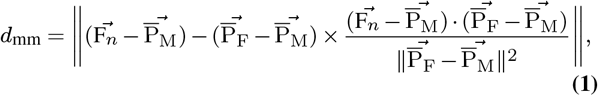

where 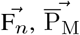, and 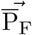 are the vectors of individual hybrid (F_n_ = F_1_ or F_2_), mean marine, and mean freshwater standardized trait values. Individuals from parent populations exhibit deviations from the line connecting parent population means due to their phenotypic variation. The average deviation did not vary among freshwater parent populations (*F*_11_,_33_._7_ = 1.45; *P* = 0.20) and accounting for parent deviation in our model does not change our conclusions (see archived analysis code). We subtracted the mean deviation observed in parents from all individual hybrid mismatch values such that hybrids with no ‘excess mismatch’ relative to parents receive a value of 0. This simply changes the intercept and has no bearing on the main conclusions which concern slopes.

Estimates of mismatch could be affected by measurement error, but measurement error is low in this dataset according to repeatability scores. Furthermore, measurement error—the absolute difference of the two ln-transformed trait measurements—is only correlated with divergence for one of fifteen traits (body depth). Freshwater population values for body depth are not significantly correlated with the phenotypic divergence between parent populations (Spearman’s rank-order correlation *P* = 0.1474). Thus, measurement error would not contribute to a ‘null’ relationship between mismatch and divergence.

We tested whether mismatch changed with the magnitude of phenotypic divergence between marine and freshwater parent populations. For simplicity, we analyze mean mismatch values for each cross type within a given marine × freshwater cross. All of our qualitative conclusions are unchanged if we analyze individual-level data using mixed models with family and population as nested random effects (see archived analysis code). Predictor variables were the (Euclidean) phenotypic divergence between the parental populations, hybrid category (F_1_ or F_2_), and their interaction.

#### Mechanisms underlying the divergence–mismatch relationship

We investigated the causal roles of dominance and phenotypic variation for generating the divergence–mismatch relationship. To determine how dominance affects mismatch for a given marine × freshwater cross, we calculated the mismatch of the mean hybrid phenotype for each population and hybrid generation. This generates a single estimate—the dominance-effect—of mismatch for each population × hybrid generation combination that results from displacement of the mean hybrid phenotype from the line connecting parent phenotypes. To determine how phenotypic variation affects mismatch, we subtracted the mismatch of the mean hybrid phenotype (the dominance-effect calculated above) from each individual’s unique mismatch value. We took the average of these differences for each family, then averaged these for a single estimate—the variance-effect—per population. We used simple linear models to test whether either the dominance- and variance-effects were associated with the magnitude of phenotypic divergence between parents; models had the dominance- or variance-effect as the response and predictor variables were as in the model testing for a divergence–mismatch relationship.

To infer why dominance and trait variation affected mismatch, we quantified general patterns of dominance and phenotypic variation hybrids. We evaluated whether traits exhibited dominance using linear mixed models where standardized trait values were the response, and the two predictor variables were (i) an additive term (fraction of the genome that is freshwater [P_M_ = 0, F_n_ = 0.5; P_F_ = 1]) and (ii) a dominance deviation (fraction of genome heterozygous [P_M_ & P_F_ = 0, F_2_ = 0.5; F_1_ = 1]) (Lynch and Walsh, 1998). Family and population were nested random effects. Dominance was only estimated for traits where the marine and freshwater parent populations were statistically distinguishable (*t*-test *P* < 0.05). To evaluate if dominance changes with the magnitude of phenotypic divergence between parents, we scaled the mean hybrid phenotype (for both F_1_s and F_2_s) such that recessive (i.e., marine-parent-like) traits had a value of 0 and dominant (i.e., freshwater-parent-like) traits had a value of 1 (transgressive values [<0 & >1] are possible); we tested if this value changed with the magnitude of phenotypic divergence between parents in mixed models with trait as a random effect. For trait variation, we calculated the variance for each trait within each family, then took the average across families. We then fit linear mixed models with mean population variance as the response, both phenotypic divergence between parent populations and hybrid category (and their interaction) as predictors, and trait as a random effect.

## Results

### Patterns of phenotypic divergence among populations

Marine × freshwater crosses differed substantially in the magnitude of phenotypic divergence between parent populations (main effect of ‘population’: *F*_1,41.8_ = 34.1; *P* < 0.0001; Fig. S2). The benthic populations from the species pairs were among the most divergent from the marine ancestor, while two highly zooplanktivorous populations that co-exist with prickly sculpin were among the least diverged (Pachena Lake and North Lake). Freshwater populations were between approximately 3–10 units diverged from the marine. The number of traits that differed significantly between the freshwater and marine parents was positively correlated with the magnitude of divergence between parent populations (Fig. S3).

### Evolution of trait mismatch in hybrids

We found support for the prediction that hybrid trait mismatch increases with the magnitude of phenotypic divergence between parents. Considering all traits together, multivariate mismatch in hybrids was positively associated with the magnitude of phenotypic divergence between parents (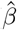 = 0.12 ± 0.028 [SE], F_1,21_ = 18.40, *P* = 0.00033) (Fig. 3). The rate of increase of mismatch with divergence (regression slope) did not differ between F_1_ and F_2_ hybrids (divergence × category interaction *P* = 0.82). Thus, for every unit of multivariate phenotypic divergence between parents, mismatch in hybrids increases by nearly 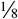 that amount. See Fig. S4 for an example of mismatch for two traits where parents have different magnitudes of divergence.

**Fig. 3.**
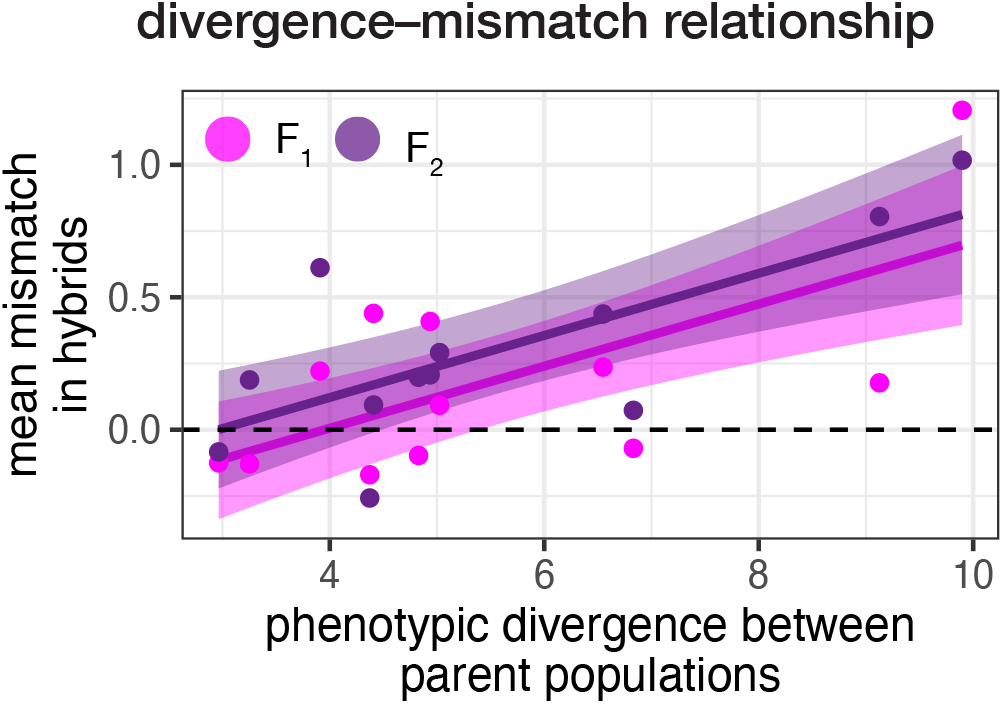
Trait mismatch in hybrids increases with the magnitude of phenotypic divergence between their parents. Each point is the mean mismatch (Eqn. 1) value across all F_1_ (pink points and lines) or F_2_ (purple points and lines) hybrids for a given marine × freshwater cross (*n* =12 per hybrid type).

### Underlying causes of the divergence–mismatch relationship

Dominance was the main cause of the divergence-mismatch relationship in F_1_ hybrids, whereas phenotypic variation was the main cause of this relationship in F_2_ hybrids (Fig. 4). Mismatch of the mean hybrid phenotype—the dominance effect—increased with the magnitude of phenotypic divergence between parents in F_1_ hybrids (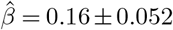; *t*_20_ = 3.09; *P* = 0.006) but not in F_2_s (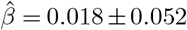; *t*_20_ = 0.34; *P* = 0.74) (Fig. 4A). Examining dominance patterns, more than ¾ (77 % or 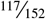) of traits were not inherited additively and 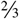 of traits exhibiting deviations from additivity tended toward recessivity (i.e., marine-like). Dominance differed among traits—for example, most F_1_ hybrids had long freshwater-like heads but also had large marine-like pectoral fins (Fig. S5). Dominance typically did not change with the magnitude of phenotypic divergence between parents, though two traits—the number of lateral armour plates and the length of pelvic spines—did show predictable evolution of dominance (Fig. S6). Returning to phenotypic variation, we found that the mismatch caused by deviation from the mean hybrid phenotype—the variation-effect—increased with the magnitude of phenotypic divergence between parents in F_2_ hybrids (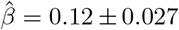; *t*_20_ = 4.69; *P* = 0.0001) but had no effect in F_1_s (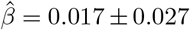; *t*_20_ = 0.64; *P* = 0.52) (Fig. 4B). Underlying this pattern was the fact that trait variation increased with phenotypic divergence between parents in F_2_ hybrids but not in F_1_s (Fig. S7). These analyses indicate that the quantitative genetic basis of the divergence-mismatch relationship differs between hybrid generations.

**Fig. 4.**
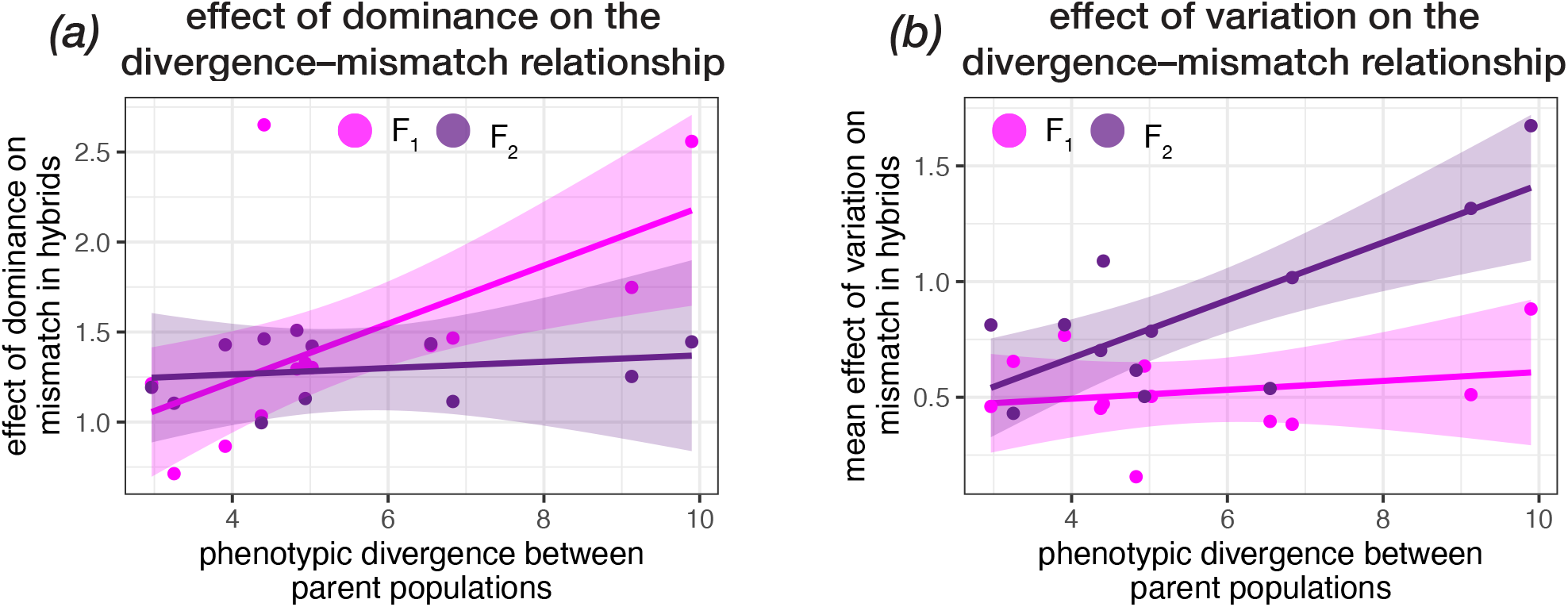
Dominance is the primary cause of the mismatch–divergence relationship in F_1_ hybrids, while variance is the primary cause of the mismatch–divergence relationship in F_2_ hybrids. Panel (A) depicts the mismatch of the mean hybrid phenotype (the *effect of dominance*), which is only caused by dominance. The slope is significantly different from zero in F_1_s but not in F_2_s. Panel (B) depicts the mean difference between mismatch values for individual hybrids and the mismatch of the mean hybrid phenotype (the *effect of variance*). The slope is significantly different from zero in F_2_s but not in F_1_s. One point is shown per marine × freshwater cross and category (i.e., F_1_ or F_2_; each *n* = 12).

## Discussion

In this study, we used experimental hybridization in stickleback to test whether the extent of trait mismatch in hybrids grows as parent populations diverge phenotypically. Trait mismatch has been associated with reduced individual fitness in recombinant stickleback (Arnegard et al., 2014) and sunflower (Thompson et al., 2021) hybrids, and a growing number of studies have used indirect inference to link mismatch in F_1_ hybrids to reproductive isolation (Cooper et al., 2018; Matsubayashi et al., 2010; Vinšálková and Gvoždík, 2007). Studies of fitness landscapes in hybrids (Keagy et al., 2016; Martin and Wainwright, 2013) and correlational selection within species (Schluter, 1994), also show patterns consistent the hypothesis that selection acts against mismatched trait combinations.

Our study was motivated by the fact that, although previous studies have documented a seemingly general relationship between ecological divergence and barriers to gene flow (Shafer and Wolf 2013), predictions of hypotheses that link adaptive divergence to the evolution of potentially maladaptive hybrid phenotypes remain untested. Comparative studies of speciation (Matute and Cooper, 2021) typically study divergence over time, whereas here we consider populations from different lakes that have diverged from a common ancestor to varying degrees in roughly the same amount of time—thus isolating the effect of phenotypic divergence. In support of our prediction, more derived and divergent freshwater parental populations tend to produce hybrids with increasingly mismatched phenotypes when each is crossed to the same marine population representing the ancestral form. The quantitative genetic underpinnings of the divergence–mismatch relationship—dominance in F_1_s and segregation variance in F_2_s—follow from theory and first principles. Below, we discuss these results in the context of speciation research and the genetics of adaptation.

### Relation to ‘intrinsic’ incompatibilities

Mismatch might be thought of as an ecological and phenotypic analogue to classic ‘intrinsic’ Bateson-Dobzhansky-Muller hybrid incompatibilities. Bateson (1909), Dobzhansky (1937), and Muller (1942) independently deduced that interactions among two or more loci likely underlie most cases of hybrid inviability and sterility, and as such ‘hybrid incompatibilities’ are explicitly defined as interactions among loci. Mismatch results from interactions among divergent loci, either via dominance or segregation, and accordingly selection against mismatched trait combinations shares this feature with ‘intrinsic’ incompatibilities. Predictions of genetically-explicit hybrid incompatibility models (Chevin et al., 2014; Simon et al., 2018) can readily be interpreted via lenses of both genetic or phenotypic mismatches, and both types of incompatibilities result in selection against opposite ancestry combinations at incompatible loci. There are some important differences between ‘intrinsic’ and ‘extrinsic’ incompatibilities, however. Intrinsic incompatibilities likely have a relatively static fitness landscape whereas the fitness consequences of mismatch are likely dynamic—with frequency, density-, and environment-dependence. Since many QTL likely underlie divergence between species (Arnegard et al., 2014; Rockman, 2012), selection against mismatch is likely highly polygenic and detecting individual loci may be exceedingly difficult except for when there are loci of major phenotypic effect.

Because mismatch increases with parental divergence, this implies that extrinsic post-zygotic isolation might evolve in a similar ‘clock-like’ manner to intrinsic post-zygotic isolation. Coyne and Orr (1989, 1997) were the first to demonstrate that reproductive isolation between populations evolves as a function of divergence time. They found that both premating and intrinsic post-zygotic isolation increased with neutral genetic distance in *Drosophila*. This work spawned a small industry (Coughlan and Matute, 2020; Matute and Cooper, 2021) reporting similar patterns in groups as diverse as orchids and fishes (Bolnick and Near, 2005; Scopece et al., 2013). The present article asks whether a potentially important predictor of extrinsic post-zygotic isolation evolves as divergence proceeds. In stickleback, genomic and phenotypic divergence are largely coincident (Miller et al., 2019; Wang, 2018), so our conclusions would likely be the same if we used genetic divergence as the predictor variable.

An important model in speciation genetics with some analogy to the results presented herein is the ‘snowball’ model of the accumulation of hybrid incompatibilities. This model, first put forward by Orr (1995), suggests that the number of hybrid incompatibilities should increase faster-than-linearly with divergence time. This is because the number of potential pairwise interactions among divergent loci—and thus pairwise incompatibilities—increases at least as fast (‘at least’ because this does not account for anything above pairwise interactions) as the square of the number of substitutions separating species. Empirical work has found support for this snowball model (Matute et al., 2010; Moyle and Nakazato, 2010; Wang et al., 2015). Our study can draw some direct parallels to the snowball model. In our data, we can determine whether a particular trait pair is mismatched if the mismatch of hybrids exceeds the typical deviation (from the axis of parental divergence) seen in parents. In this analysis, we find evidence of a snowball in F_1_ hybrids but not in F_2_s where the increase is linear (see Fig. S8). We view this analysis as purely heuristic because trait pairs are not independent, though this issue also affects other studies of incompatibilities. Nevertheless, trait mismatches do seem to snowball in a similar manner to intrinsic incompatibilities. The fitness consequences of this snowballing, however, are unclear, and depend fundamentally on how the fitness-effects of incompatibilities interact when present in the same genetic background (Guerrero et al., 2017).

### Dominance of alleles used for adaptation to freshwater

We found that derived traits are typically not dominant in stickleback. Although this unexpected if adaptation were from new mutation (Haldane, 1927), this finding is consistent with what is known about adaptation to freshwater habitats in stickleback proceeding predominantly via the reassortment of existing standing genetic variation (Jones et al., 2012b; Nelson and Cresko, 2018). Specifically, theory predicts that dominance does not affect the fixation probability of alleles in the standing variation (Orr and Betancourt, 2001) and that their fixation probability increases with their frequency in the population (MacPherson and Nuismer, 2017). We also found that deviations from additivity were in a recessive direction (i.e., toward marine) more often than in a dominant direction. These results are consistent with the findings of Miller et al. (2014), who used QTL mapping to measure dominance in an F_2_ marine (from Japan) × freshwater (Paxton Lake benthic) cross, and found that most QTL were additive or partially additive with a slight but significant bias toward recessivity. Why might most QTL that contribute to freshwater adaptation be recessive? Theory and empirical studies indicate that deleterious alleles are maintained at higher frequencies when they are recessive (Charlesworth and Hughes, 1998; Robinson et al., 2018; Simmons and Crow, 1977; Willis, 1999), and if freshwater-adaptive alleles are deleterious in the sea this could explain why there is a slight bias toward recessive QTL in the sea.

With two notable exceptions, dominance did not change predictably with the magnitude of phenotypic divergence between parents. This finding is not surprising because the mean dominance coefficient of alleles fixed (or that rise to high frequency) from standing variation should not depend on the distance of a phenotypic optimum. In stickleback, even highly derived freshwater populations rely on standing variation for adaptation (Jones et al., 2012a; Nelson and Cresko, 2018). However, dominance of two traits—the number of lateral armour plates and the length of the pelvic spine—varied predictably with the magnitude of phenotype divergence between parent populations. Plate number was recessive in crosses with ancestor-like freshwater populations and increasingly dominant when the marine was crossed with more derived benthic-feeding populations. Plate reduction is known to be largely caused by a single large-effect variant at the *Eda* locus (Archambeault et al. 2020; Colosimo et al. 2005), which is fixed for the freshwater allele in all but one (North Lake) of the freshwater populations considered here. Previous studies have shown that the alleles that reduce plate number do themselves modify the dominance of *Eda* (Colosimo et al., 2004). Our result adds to the understanding of this trait by showing that the net effect of alleles that modify dominance of lateral armour plates is predictable based on the phenotype of the freshwater parent. We also found that the length of pelvic spines was largely dominant in crosses with an ancestor-like freshwater population and largely recessive in crosses involving derived populations. The pelvic girdle, the presence and size of which is governed by *PitX1* (Chan et al., 2010), is also known to be recessive in pelvis-absent populations (i.e., it is present when heterozygous and only absent when homozygous for the freshwater allele). Little is known about dominance modifiers of *PitX1*, and this would be a valuable topic for future work. Thus, dominance of most traits does not evolve predictably. Although hypotheses about the evolution of dominance abound (Fisher, 1928; Wright, 1934), we cannot determine here whether the evolution of dominance for two traits documented here is incidental or adaptive.

### Caveats, future directions, and conclusions

The biggest and clearest limitation of our study is its lack of a direct link between mismatch and fitness. In stickleback specifically, selection against mismatched trait combinations has been shown twice (Arnegard et al., 2014; Keagy et al., 2016), but only in crosses between benthic and limnetic populations. Selection against natural marine × freshwater hybrids has been inferred in hybrid zones from the steepness of clines (Vines et al., 2016), as well as observations of heterozygote deficit and cytonuclear disequilibrium (Jones et al., 2006). In the Little Campbell River studied here, Hagen (1967) inferred that selection against marine × freshwater hybrids is “very intense”, although the specific mechanisms of selection were unclear. Clearly, we must begin to conduct comparative studies where mismatch can be linked to fitness directly. In hybrid zones, biologists can estimate the strength of selection against hybrids by, among other methods, measuring cline width and back-crossing rates. Future studies could leverage areas of ongoing hybridization to evaluate whether phenotypic mismatch measured in crosses predicts the strength of selection against natural hybrids (Barton and Hewitt, 1985; Harrison, 1993). Experimental arrays with recombinant hybrids and parents could be used to relate mismatch to back-crossing rates to identify its effectiveness as a barrier to gene flow. Experimental evolution studies could also be used to robustly estimate the generality of the divergence-mismatch relationship and its effect on hybrid fitness. It would be particularly valuable to identify generalities about whether the mismatch-fitness relationship is linear, diminishing, or accelerating. Clearly, more work is necessary to solidify our general understanding of the fitness effects of mismatch.

Some of our findings are more likely to be general than others. In particular, our results regarding F_2_ hybrid phenotypic variation increasing with the magnitude of divergence between parents were predicted from theory (Barton, 2001; Chevin et al., 2014; Slatkin and Lande, 1994). It is therefore a reasonable expectation that this particular finding might extend to other systems. Because dominance is somewhat idiosyncratic, it is less clear how general our finding that dominance causes a divergence-mismatch relationship in F_1_s will be, though dominance in hybrids is the rule rather than the exception (Thompson et al., 2021). Since the divergence–mismatch relationship we document here might be an important mechanism driving progress toward speciation, establishing which aspects are idiosyncratic and which are general seems worth the effort.

Ultimately, there are many causes of speciation and trait mismatch will be one of many proximate causes of reproductive isolation. Any mismatch would likely operate alongside other well-documented processes such as assortative mating (Coyne and Orr, 2004; Jiang et al., 2013; Rundle et al., 2000) and selection against immigrants (Nosil et al., 2005). Empirical estimates of the relationship between ecological divergence and hybrid fitness (Edmands 1999; Funk et al. 2006; Shafer and Wolf 2013), or neutral divergence and hybrid fitness (Coughlan and Matute, 2020; Matute and Cooper, 2021), invariably find that these relationships are noisy. Because F_1_ hybrid mismatch is prevalent in many systems (Thompson et al., 2021), it could be an immediate and powerful barrier to gene flow between many diverging lineages. As shown above, it might even ‘snowball’. Future studies clarifying the importance of mismatch for speciation are sorely needed.

## Author contributions

KAT and DS designed the study. KAT conducted fieldwork, made crosses, and collected samples. KAT and AKC raised animals, collected data, and co-wrote the first draft of the manuscript. AKC and KAT analyzed the data with input from DS. All authors revised the manuscript.

## Data accessibility

All data and analysis code used in this article will be deposited in a permanent repository (e.g., Dryad) following publication. They are available to reviewers of the submitted version of the manuscript.

## Acknowledgements

Feedback from D. Irwin, L. Rieseberg, S. Otto, the Schluter Lab (UBC), and the Schumer Lab (Stanford) improved the manuscript. L. Alford, M. Andersen, M. Ankenmann, J. Bizon, S. Blain, L. Chavarie, A. Chhina, K. Chu, J. Dafoe, N. Frasson, C. Gerlinsky, A. Jevtic, M. Kinney, S. Larter, M. Mikkelsen, K. Nikiforuk, A. Munzur, M. Osmond, M. Roesti, J. Rolland, G. Singh-Varma, M. Urquhart-Cronish, and J. Viliunas provided logistical support in the field and/or lab. Jim & Arron kindly provided accommodation on Nelson Island. Ralf Yorque shepherded the authors through all aspects of the project. B. Gillespie, E. Lotto, and P. Tamkee provided infrastructure support. We are grateful to the Semiahmoo First Nation, in particular J. Charles and J. Cook, for facilitating our collections from the Little Campbell River. AKC was supported by an NSERC USRA. KAT was supported by an NSERC Canada Graduate Scholarship, a UBC Four-Year Fellowship, and a British Columbia Graduate Fellowship. DS was supported by an NSERC Discovery Grant, Genome Canada, Genome BC, a Canada Research Chair (Tier I), and a UBC Killam Professorship. R. Henriques created the 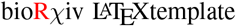. The University of British Columbia is located on the traditional and unceded territory of the Musqueum First Nation.

## Supplementary material

### A. Supplementary methods

#### A.1. Details of crossing protocol and fish husbandry

Female stickleback were selected for spawning when their abdomens were sharply angled at the cloaca and the first egg was visible. We gently squeezed the sides of the female fish’s body to release the eggs into a Petri dish containing water from her source habitat (tank, lake, or river water). Mature male stickleback were identified by their bright blue colouration and red throat. Male fish were euthanized with an overdose of MS-222, and then testes were extracted from the body cavity using fine forceps after making a small incision beginning at the cloaca. We used a small paintbrush to release sperm from testes and to ensure that sperm contacted all eggs. The live fish and fertilized clutches were transported to the InSEAS aquatic facility at the University of British Columbia, Vancouver, British Columbia, Canada.

All fish were hatched in 100 L aquaria with room temperature between 17 and 19 °C and a photoperiod that followed local dawn and dusk times. Instant Ocean^®^ Sea Salt was added to maintain a salinity of 5 ppt in all tanks. Fry were fed live brine shrimp nauplii. Chopped frozen bloodworms were added to the diet when fish were large enough, and then finally adult-sized fish were fed full size frozen bloodworms and frozen mysis shrimp *ad libitum* (Hikari Bio-Pure^®^).

We sampled fish for phenotype measurements typically when the mean standard length of a family was approximately 40 mm. Sticklebacks have adult morphology at this stage and are not sexually reproducing. Due to occasional logistical constraints, some tanks were sampled at earlier or later mean standard length sizes. Also, due to logistical constraint, all populations except for Paxton benthic and Paxton limnetic were collected in 2017.

#### A.2. Repeatability

We evaluated the repeatability of our measurements to determine the fraction of variance that could be attributable to measurement error. Repeat measurements were made on at least 25 fish. For linear measurements (including pectoral fin length), we took two separate photographs of each fish (or fin) and made the repeated measurements on these separate photographs. Second photographs were made after returning and then removing the fish (or fin) from its storage vial. Count and gill raker measurements were made on the original specimens. In all cases except pectoral fin length, first and second measurements were made more than one year apart.

#### A.3. Data diagnostics

We checked for outliers in the raw data and evaluated outlier individuals to ensure they were not caused by measurement or transcription error. If fish were inadvertently measured twice, we averaged trait values across measurements. Fish with broken second dorsal spines were removed from the dataset. One fish was removed because it had an unusual body shape—qualitatively appearing as if it had failed to inflate its swim bladder—and it was an extreme outlier in Normal Q-Q plots and in standardized residuals vs. leverage plots. Such phenotypes seem to be caused by environmental factors (e.g., a too-powerful air stone) rather than biological factors (e.g., hybrid incompatibilities).

### B. Pairwise trait mismatch

For pairwise mismatch metrics, separate regressions were carried out for each pair of traits. We evaluated the statistical significance of the regression models, as well as the distribution of regression coefficients across models to generate inferences about the relationship between adaptive divergence between parents and mismatch in hybrids.

We examined the relationship between phenotypic divergence and hybrid mismatch for pairs of traits at a time. We found that pairwise trait mismatch was significantly associated with parent trait divergence for 25 of 105 trait pairs in F_1_s, and in F_2_ hybrids this was 32 of 105 trait pairs (37 %). All significant slopes were positive (Fig. S9). The mean absolute slope of significant relationships was approximately 0.08 in both hybrid types. Significant pairwise regressions typically involved one or both of: pelvic spine length, pelvic girdle length, or lateral plate count, but several divergence-pairwise mismatch regressions lacking those traits were significant (see archived analysis code).

### C. Supplementary figures

**Fig. S1.**
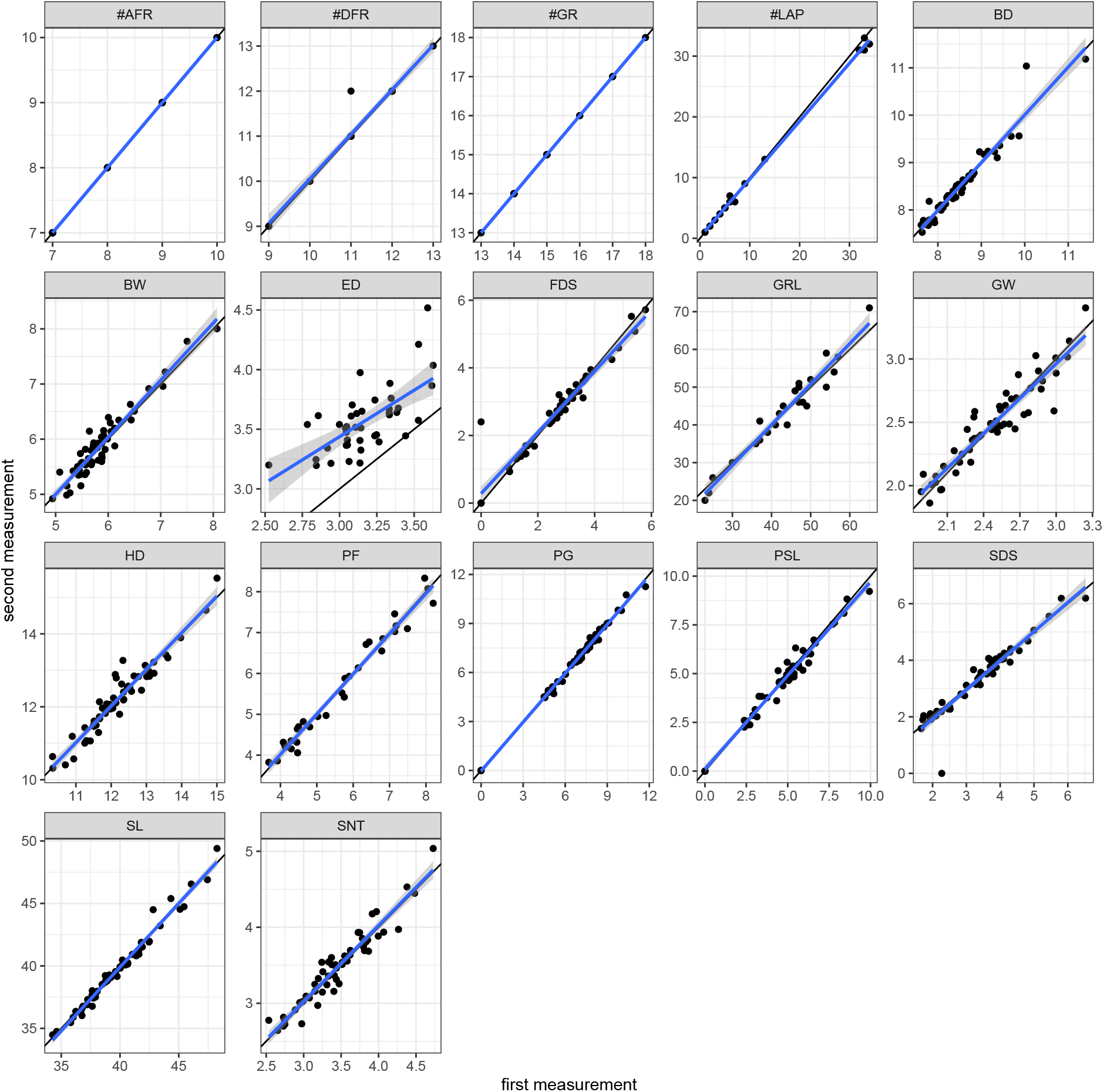
Repeatability data for all measured traits. All plots show the first and second measurements made on all traits. Black lines are 1:1 lines, and blue lines are linear regressions. Trait codes are as in Fig. 2B. The fish with a value of ‘0’ for second dorsal spine (SDS) likely had its spine broken off during the time-frame between first and second measurements. All *r*_Pearson_ > 0.9, except for eye diameter (ED), which was dropped from the analysis.

**Fig. S2.**
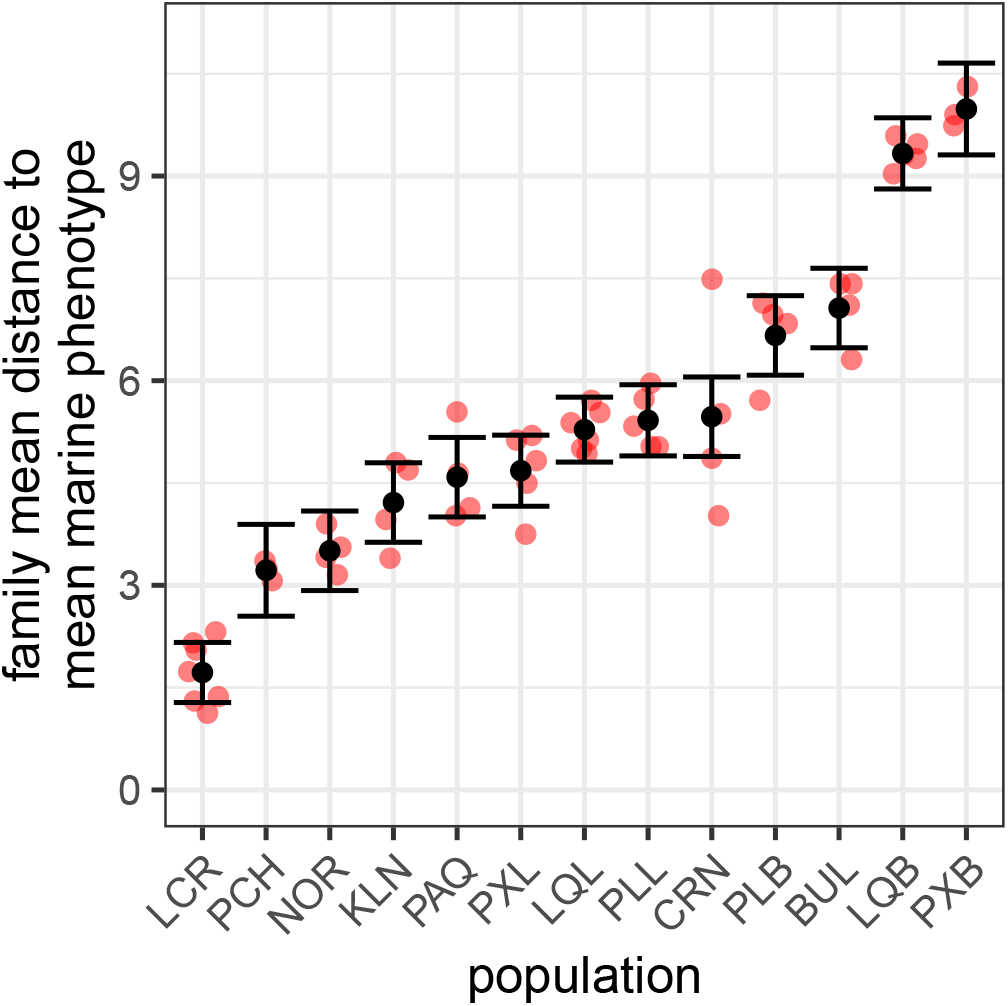
Among-population variation in the phenotypic divergence to the marine ancestor. Red points show the phenotypic distance from each family’s vector of trait means to the mean marine phenotype. Black points and lines are means and 95 % confidence intervals extracted from the model using `visreg` (Breheny and Burchett, 2017). Population codes (left to right): LCR—Little Campbell River; PCH—Pachena Lake; NOR—North Lake; KLN—Klein Lake; PAQ—Paq (Lily) Lake; PXL—Paxton Lake Limnetic; LQL—Little Quarry Lake Limnetic; PLL—Priest Lake Limnetic; CRN—Cranby Lake; PLB—Priest Lake Benthic; BUL—Bullock Lake; LQB—Little Quarry Lake Benthic; PXB—Paxton Lake Benthic. The Little Campbell River population is not zero because family means are not identical to the mean of family means.

**Fig. S3.**
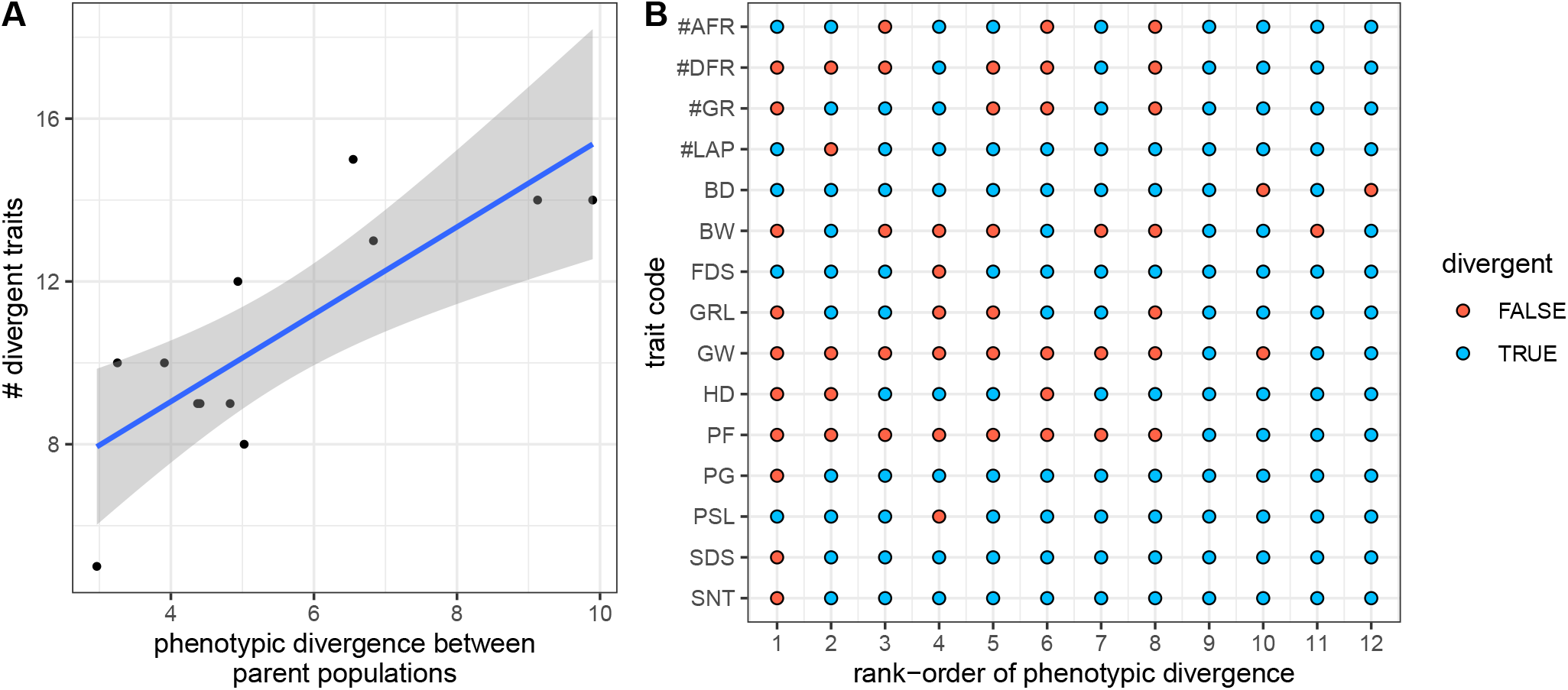
Phenotypic divergence between parents is positively associated with the number of traits that differ between them. (A) Each point is one freshwater population. The number of traits is the number that differ significantly between the freshwater population and the anadromous ancestor according to a Bonferroni-corrected *t*-test. For every unit of multivariate phenotypic divergence, there is one additional divergent trait (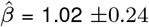; *F*_1,10_ = 18.4; *P* = 0.0016). Panel (B) shows whether or not individual traits (codes as in Fig. 2B) differ between the freshwater and marine parent for all twelve populations set out on the *x*-axis by rank-order of phenoypic divergence (see Fig. S2 for rank order).

**Fig. S4.**
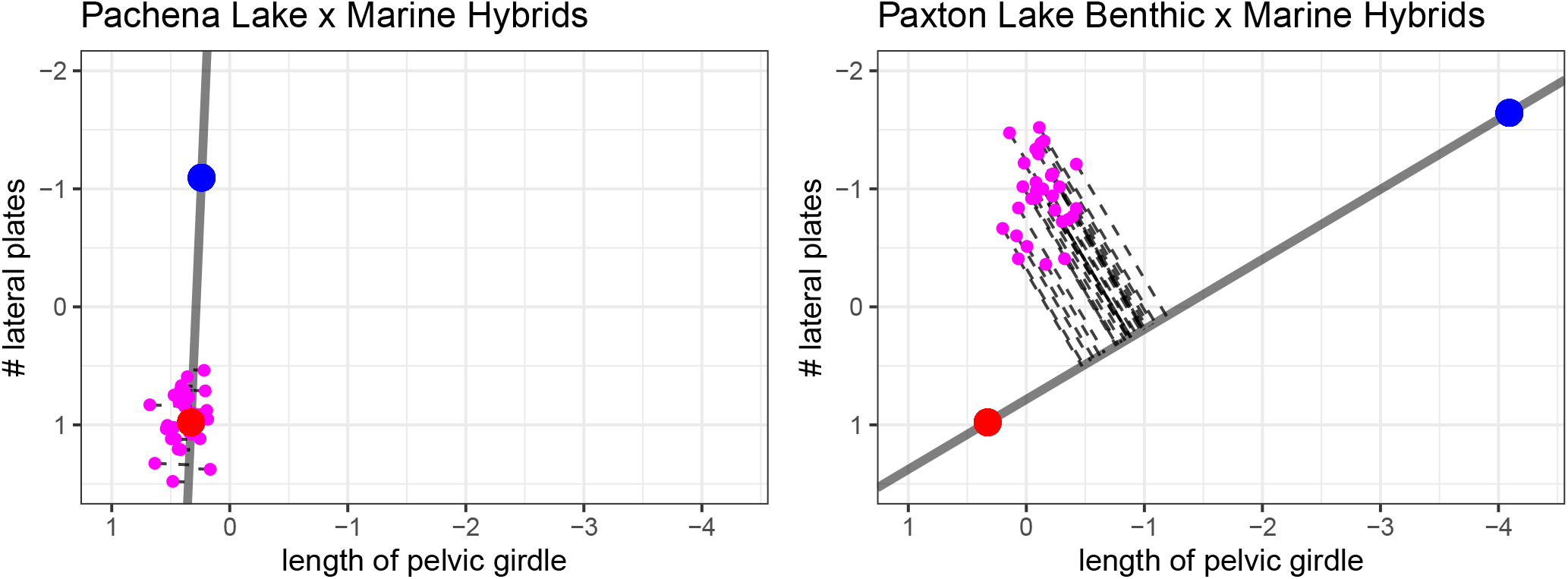
Visualization of pairwise mismatch for two traits using our empirical trait data. Each plot shows the scaled phenotype data used in main text analyses. The red points indicate the mean of the freshwater parent population and the blue points indicate the mean of the marine parent population. Mismatch is the length of the perpendicular line connecting black points—individual F_1_ hybrids— to the line connecting parent mean phenotypes. The higher average mismatch of the Paxton benthic hybrids (*μ* = 1.54) than Pachena Lake hybrids (*μ* = 0.087) results from opposing dominance: the pelvic girdle phenotype resembles the marine ancestor whereas the plate number is similar to the freshwater parent.

**Fig. S5.**
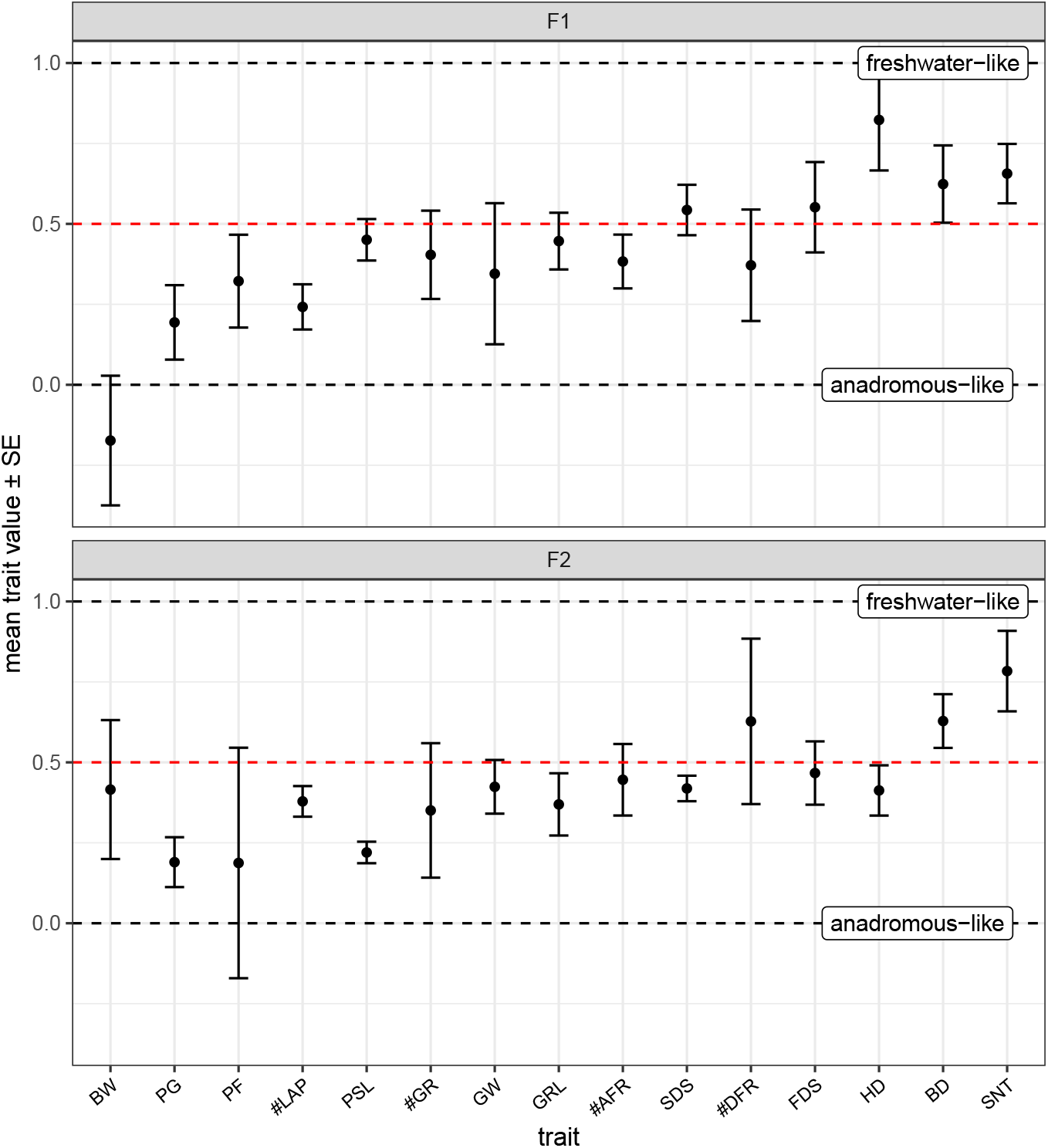
Dominance of the freshwater phenotype in hybrids for different traits. Trait values are scaled such that populations complete recessivity (i.e., marine-parent-like) is a value of 0, additivity is a value of 0.5, and complete dominance (i.e., freshwaterparent-like) is a value of 1. Points depict the mean (± SE) *scaled* trait value—calculated across all populations—for all measured traits (*n* = 862 fish including F_1_ hybrids and parents). The dashed lines at 0 and 1 represent the ancestral marine parent and derived freshwater parent trait values, respectively, and the red dashed line at 0.5 represents the mid-parent value.

**Fig. S6.**
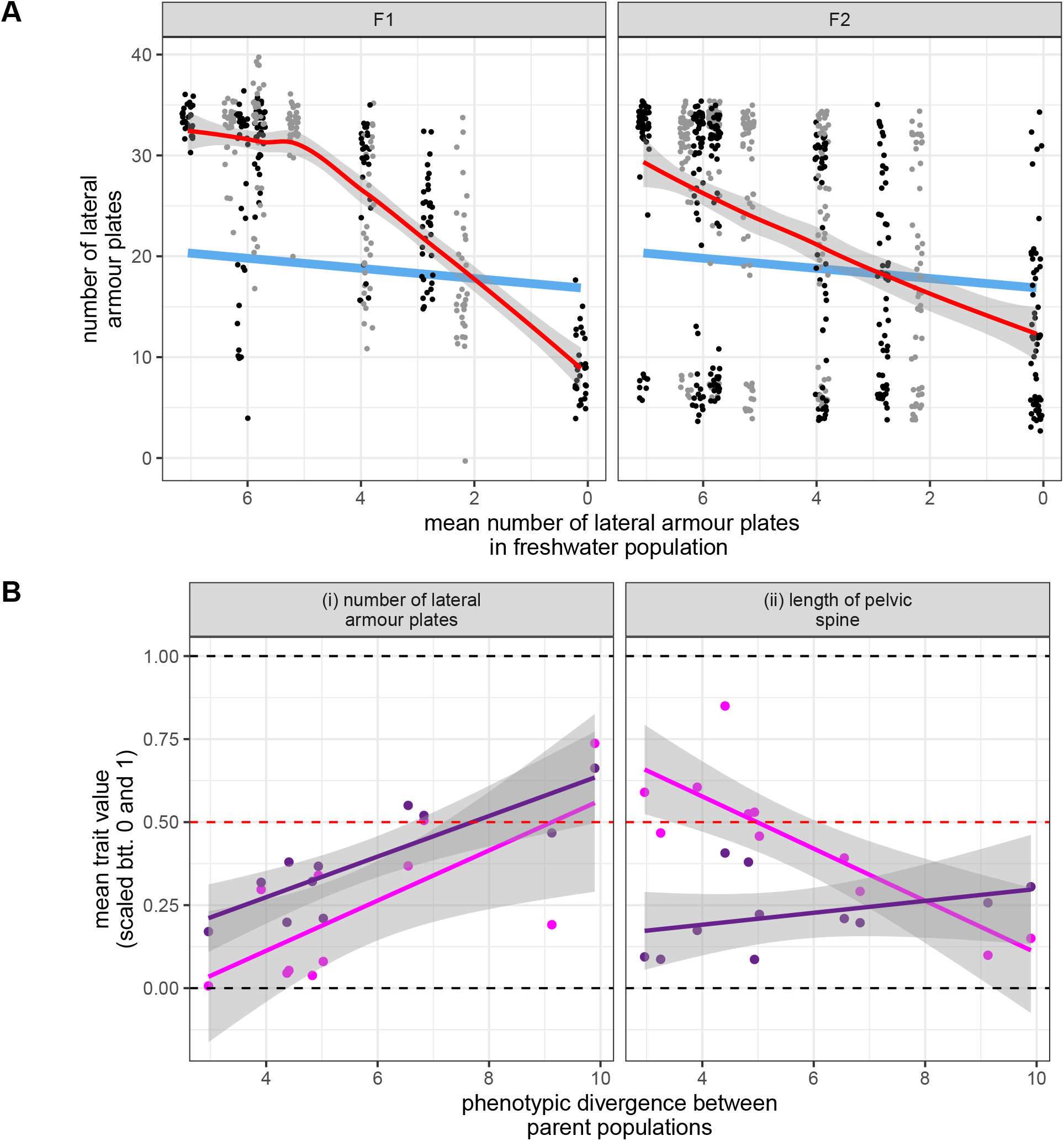
Evolution of dominance. Panel (A) shows individual values for lateral plate counts for both F_1_ (left) and F_2_ (right) hybrids. The red line is a loess-smooth fit to the data, and the blue line is fit to the parental midpoint. The *x*-axis is reversed and more derived populations are on the right. Grey and black coloured points simply demarcate populations with adjacent mean divergence values (as in chromosomes on a Manhattan plot). In both hybrid types, dominance evolves towards less recessivity as divergence proceeds. Panel (B) shows scaled trait values (as in Fig. S5) for F_1_ (pink) and F_2_ (purple) hybrids for both armour plates (left) and pelvic pine length (right). Panels A and B(i) show the same data plotted in different ways (we do not show the raw data for pelvic spine length since the high variability renders visualization challenging). Pelvic spines are increasingly recessive as divergence proceeds in F_1_ hybrids, but not F_2_s.

**Fig. S7.**
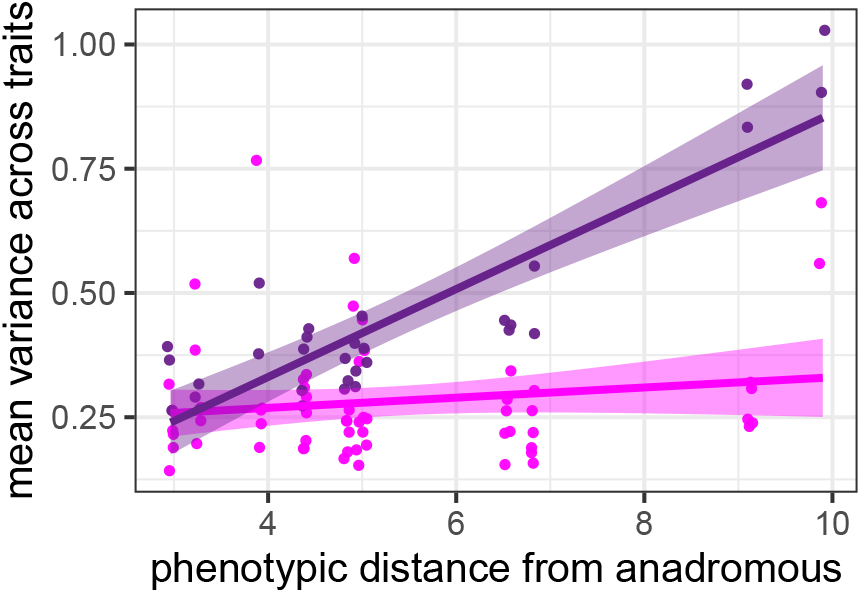
Phenotypic variation increases with the magnitude of phenotypic divergence between parents in F_2_ (purple) hybrids but not in F_1_ hybrids (pink). Points represent the mean of variances across all 15 traits within each independent family (minimum *n* = 5). F_1_ and F_2_ hybrids are distinguished by colour. Linear measurements are ln-transformed, so this result is not simply due to scaling means and variances (count data are raw but values are *lower* in more derived crosses for all meristic traits). Points are horizontally jittered to ease visualization.

**Fig. S8.**
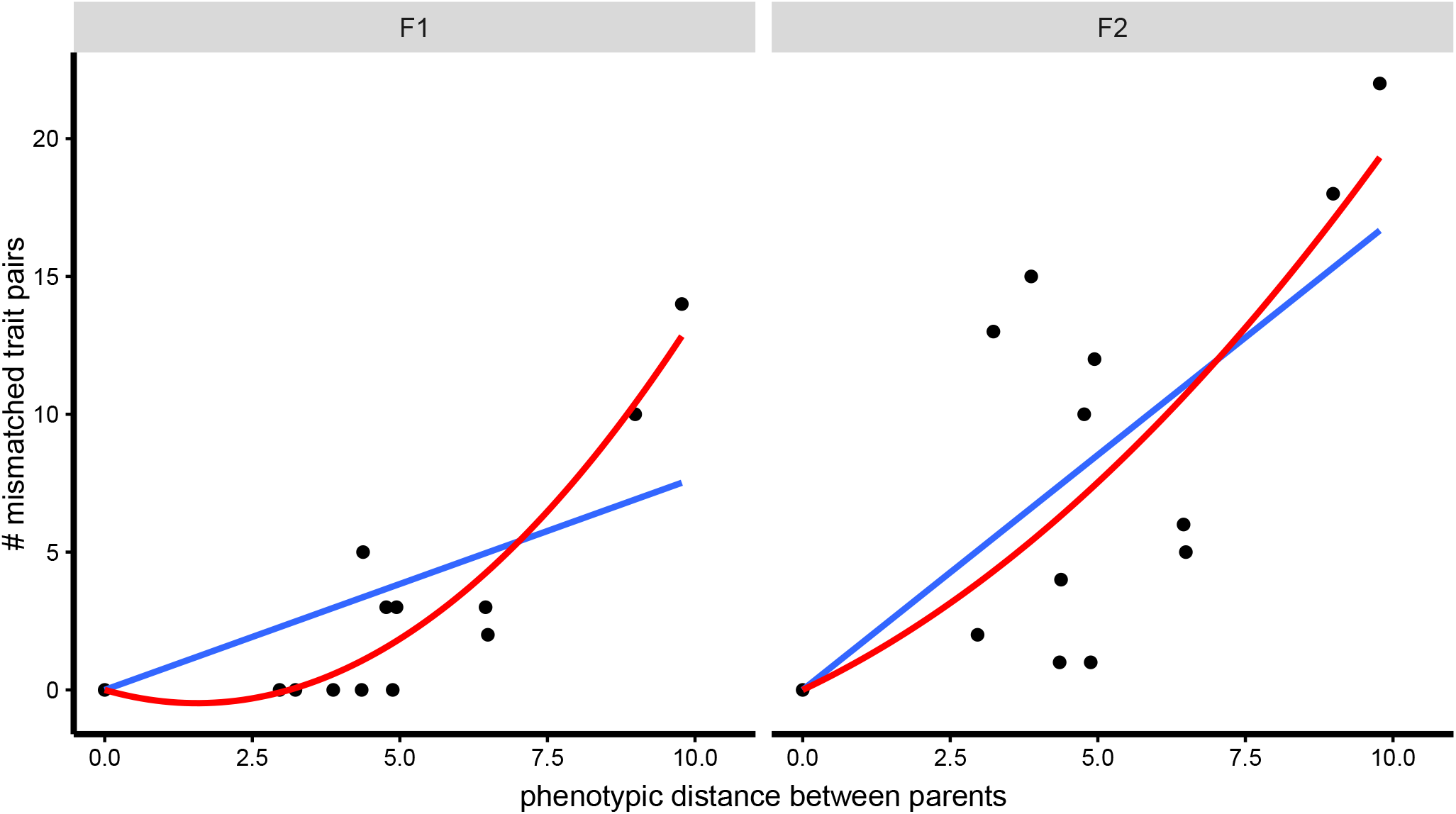
The number of mismatched trait pairs ‘snowballs’ with the magnitude of phenotypic divergence between parents, but only in F_1_ hybrids. The y-axis shows the number of trait pairs with significant mismatch, determined using *t*-tests which tested the null hypothesis that the difference between hybrid mismatch (with weighted pooling by families) and the pooled ‘mismatch’ across pure freshwater populations was 0. *P*-values were Bonferroni-corrected. The plot and regression lines are modelled after the ‘snowball’ studies of Moyle and Nakazato (2010) and Matute et al. (2010). The blue lines are linear regressions and the red lines are quadratics. Results hold if the intercept is not forced through zero, and if the ‘origin’ datum is omitted.

**Fig. S9.**
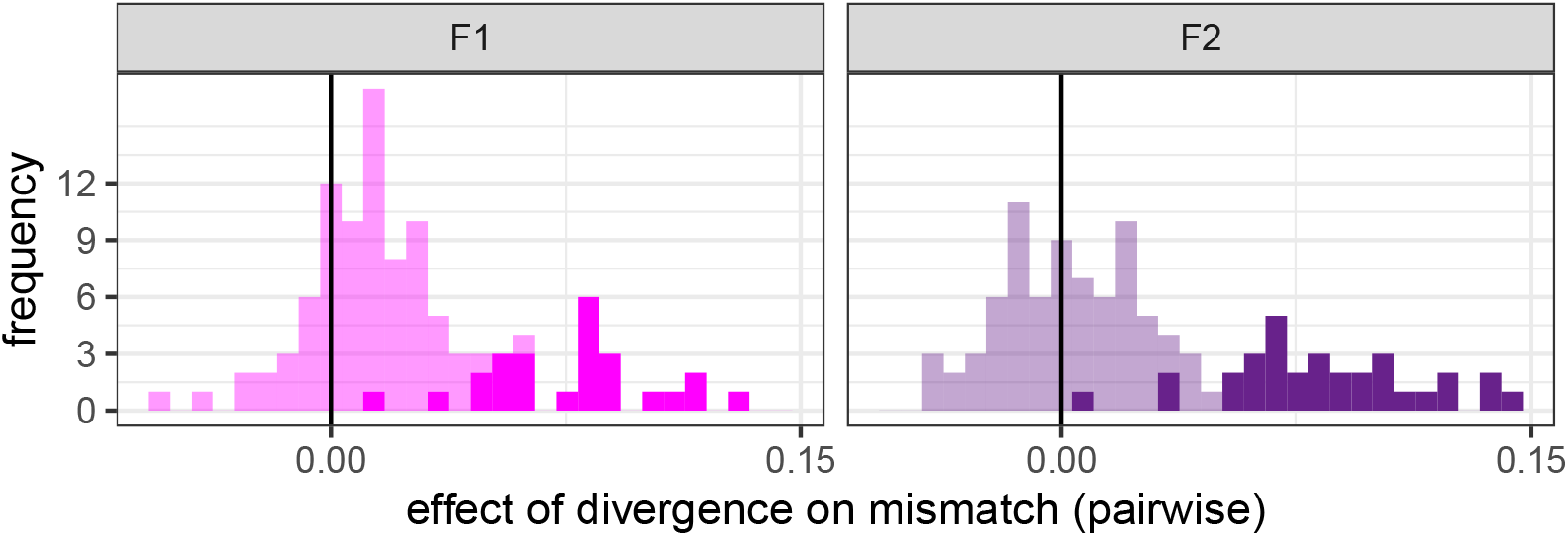
Distribution of divergence-mismatch slopes 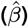 for pairwise analyses. We are showing the frequency distribution of slopes for the pairwise mismatch analyses in F_1_ and F_2_ hybrids. These slopes capture how mismatch for a pair of traits changes with the magnitude of phenotypic divergence between parents for those two traits. Significant slopes (*) are shown in a darker shade than non-significant (n.s.) slopes.

